# Dissociation of connectivity for syntactic irregularity and perceptual ambiguity in musical chord stimuli

**DOI:** 10.1101/2020.01.12.903583

**Authors:** Chan Hee Kim, Seung-Hyun Jin, June Sic Kim, Youn Kim, Suk Won Yi, Chun Kee Chung

## Abstract

Previously syntactic irregularity has been most studied with chord sequences. However, the same chord may be interpreted as having different harmonic functions, implying perceptual ambiguity. Hence, syntactic irregularity and perceptual ambiguity may be processed simultaneously. We devised 3 different 5-chord sequences in which the ending chord differed with the tonic (T), submediant (SM), and supertonic (ST). In terms of syntactic regularity, T is most regular, ST is most irregular. However, in terms of perceptual ambiguity, the most irregular ST had the salient highest voice. Therefore, the SM was the most ambiguous condition. We investigated how the human brain separates syntactic irregularity and perceptual ambiguity in terms of effective connectivity in bilateral inferior frontal gyri (IFGs) and superior temporal gyri (STGs) with magnetoencephalography in 19 subjects. Correct rate was lower for the most ambiguous chord (SM) (*P* = 0.020) as expected. Connectivity from the right to the left IFG was enhanced for the most irregular chord (ST) (*P* = 0.024, false discovery rate (FDR) corrected), whereas connectivity from the right to the left STG was enhanced for the most ambiguous chord (SM) (*P* < 0.001, FDR corrected). The correct rate was negatively correlated with connectivity in the STG, further reflecting perceptual ambiguity (*P* = 0.026). We found that syntactic irregularity and perceptual ambiguity in music are dissociated in connectivity between bilateral IFGs and STGs, respectively.

**Significance Statement:** We provide the first neurophysiological evidence of the processing of perceptual ambiguity, other than syntactic irregularity, implied in musical chords. We found that the notion of “perceptually ambiguity” is applicable to musical chord stimuli different in syntactic irregularity, and that perceptual ambiguity is separate from syntactic irregularity. Our data demonstrate that the brain interprets the three conditions of musical chords as both “from regular to irregular” and “from ambiguous to unambiguous” conditions simultaneously. This study is the first to unveil dissociation of connectivity by syntactic irregularity and perceptual ambiguity involved in musical chord stimuli.

## Introduction

Previous studies on musical syntax using chord sequences have only focused on classifying syntactic regularity and irregularity of harmony using chord sequences (Maess et al., 2001; Koelsch et al., 2002a; Patel, 2003; Kim et al., 2011). However, the same notes, intervals, or chords can be interpreted as having different harmonic functions, implying perceptual ambiguity. Even composers often exploit such potential ambiguities. Hence, the two processes of syntactic irregularity and perceptual ambiguity are overlapping.

The term “ambiguity” generally refers to uncertainty caused by two or more interpretations of an object, which is distinct from “vagueness” referring to interpretive uncertainty implied in a unity of different meanings (Tuggy, 1993; Kennedy, 2019). Ambiguity can occur in music, too, as in language. It occurs on various levels. For instance, metric ambiguity can happen when two or more possible metric interpretations are possible such as hemiola between duple and triple meter (Agawu, 1994). In tonal music, the same triad of C-E-G can be interpreted as the tonic (I) of C major or the dominant (V) of F major. Tritone substitution between the V7 chord and the flat II7 chord often occurs in jazz as the tritone (B-F), for instance, can be either a part of the chord of G7 or that of Db7 (Karpinski, 2012).

Since most previous music studies used 2 different chord sequences, syntactic irregularity and perceptual ambiguity could not be separated. However, if we use 3 different chord sequences, syntactic irregularity and perceptual ambiguity could not be in the same direction. We found the notion of ambiguity applicable to the level of cadences in music. Figure 1A shows three 5-chord sequences with different endings on the three chords of tonic (T), submediant (SM), and supertonic (ST), which are different in syntactic irregularity (T = most regular, SM = less regular, ST = most irregular).

**Figure 1.**
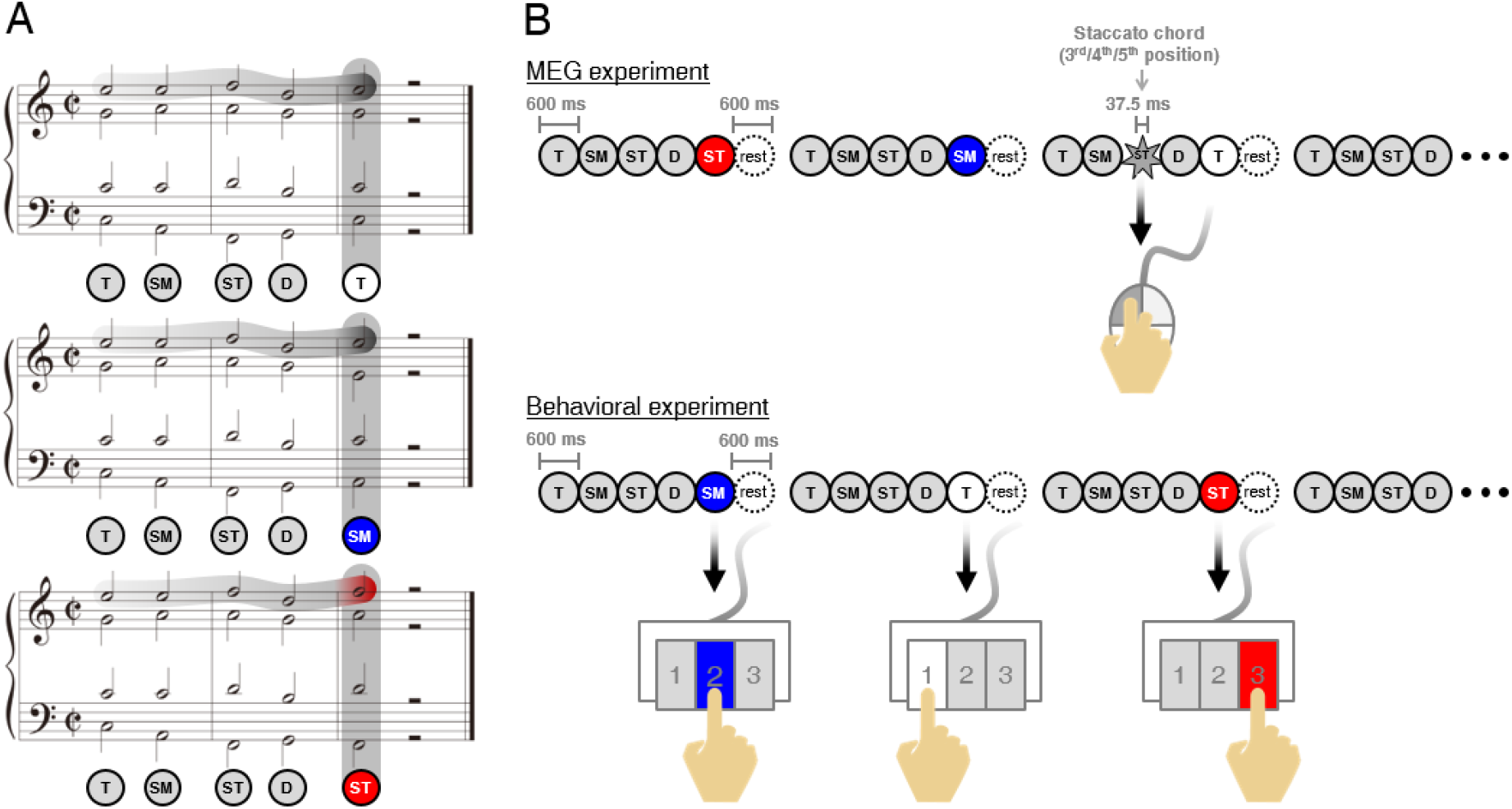
Musical stimuli and experimental paradigm. (A) Five chords in each condition differ in harmony and melodic line of the ending chords following dominant (D). Harmonies are tonic (T), submediant (SM), and supertonic (ST). The highest voices of melodic line are “E” (dark gray shadow) in the T and SM, an “F” (red shadow) in the ST. (B) In the MEG experiment (top), the participants listened to the three conditions carefully and were asked to detect the staccato chord among chord sequences in the three conditions and to click a mouse (to check the level of attending the conditions). The staccato sequences (10% of all sequences) were excluded in the MEG data analysis. In the behavioral experiment (bottom), the participants discriminated the three conditions and responded by using the 1, 2, and 3 buttons on the keypad.

The SM was the most ambiguous condition among the three conditions. The SM is difficult to distinguish from the T, because it is a substitute chord of the T, sharing the same notes of “C” and “E”. It is a deceptive cadence implying both of closure and progression, and it has the non-salient melodic line of “E” which is the same with the T. The SM is also difficult to distinguish from the ST, because the sonic quality is the same minor triad as the ST and both SM and ST are syntactically irregular chords, compared to the T. Given these relations, the SM is most ambiguous, and the ST is most unambiguous.

We investigated how perceptually ambiguous chord is processed in the brain, and how the human brain separates syntactic irregularity and perceptual ambiguity in terms of effective connectivity. Using linearized time delayed mutual information (LTDMI; see Materials and Methods section) (Jin et al., 2010), we estimated connectivities modulated by the most irregular and ambiguous conditions among twelve directional connectivities between the bilateral inferior frontal gyri (IFGs) and superior temporal gyri (STGs).

## Materials and Methods

### Ethics statement

In the present study, we used the same data sets of our previous study (Kim et al., 2019) that was approved by the Institutional Review Board of the Clinical Research Institute, Seoul National University Hospital (H-1001-020-306), and applied novel hypotheses and analyses. All participants provided informed consent of written form prior to the experiments.

### Participants

All participants were 19 females (mean age, 24.3 ± 3.0 years) of 9 music-majors and 10 non-music-majors. They were all had normal hearing and were right-handed.

### Musical stimuli

We used 3 different conditions: T, SM, and ST (Fig. 1A). The T was comprised of “tonic-submediant-supertonic-dominant-tonic”. The ending chord of “tonic” was replaced with “submediant” and “supertonic” in the SM and ST, respectively. The chords from first to fourth were the same in all conditions. The “F” of the highest voice in the final chord in the ST was different from “E” in the other two conditions. In each condition, the duration of a chord was 600 ms. A chord sequence totaled 3600 ms, including five chords and a 600 ms resting period (Fig. 1B). All conditions transposed into 12 major keys, were randomly shuffled in each session and were recorded at 100 BPM using Cubase 5 (Steinberg Media Technologies, Hamburg, Germany) software. The intensity was normalized in each wave file (sampling rate: 44.1 KHz; 16-bit; stereo; windows PCM) using Cool Edit Pro 2.1 (Syntrillium Software Corporation, Phoenix, AZ, USA). The piano timbre (Bösendorfer 290 Imperial grand) in each chord was created by Grand 3 (Steinberg Media Technologies, Hamburg, Germany) software.

### MEG recording

The whole experimental paradigm was comprised of 3 behavioral test sessions after 6 MEG recording sessions (Fig. 1B). Each MEG session included 100 sequences consisting of 30 sequences per condition and 10 staccato sequences. In individual staccato sequences, a staccato chord of 37.5 ms duration was presented in the third, fourth, or fifth chords. The participants were asked to detect staccato chord and to respond using a mouse. The response for staccato sequences was excluded in the MEG data analysis. In each behavioral session after MEG recording, 12 sequences per condition were randomly presented. All participants were asked to identify each condition of T, SM, and ST using the 1, 2, and 3 buttons on keypad. The musical stimuli were presented at the sound pressure level of 65 dB into MEG-compatible tubal insert earphones (Tip-300, Nicolet, Madison, WI, USA) using the STIM^2^ (Neuroscan, Charlotte, NC, USA) system. The whole experiment took about two hours. MEG signals were recorded in a magnetically shielded room using a 306-channel whole-head MEG System (Elekta NeuroMag VectorView™, Helsinki, Finland), with a sampling rate of 600.615 Hz using 0.1–200 Hz band pass filter. Electrooculograms (EOG) and electrocardiograms (ECG) were simultaneously recorded to later remove ocular and cardiac noise.

### MEG analysis

The environmental magnetic noise of raw MEG signals was eliminated by the temporal Signal-Space Separation (tSSS) algorithm in MaxFilter 2.1.13 (Elekta Neuromag Oy, Helsinki, Finland) (Taulu and Simola, 2006; Taulu and Hari, 2009). The 204 orthogonal planar gradiometer in 102 locations was used in the further analysis procedure.

Source analysis of four ROIs (bilateral IFGs and STGs) was performed using BESA 5.1.8.10 (MEGIS Software GmbH, Gräfelfing, Germany). The multiple equivalent current dipoles (ECDs) were estimated with the same procedures as on our previous studies (Kim et al., 2011; Kim et al., 2014). After the ECDs of P2’s magnetic counterpart (P2m) were estimated in the peak latency of 180–190 ms for the first “tonic” chord of the chord sequence in the bilateral STGs, the ECDs of ERANm were estimated in 140–220 ms for all ending chords (mean of the tonic, submediant, and supertonic chords) in the bilateral IFGs. The multiple dipoles were more than 80% of the goodness of fit (GOF). The estimated dipoles in the IFG were superior and anterior to these in the STG (Maess et al., 2001; Kim et al., 2011; Kim et al., 2014). The x, y, and *z* in Talairach coordinates (millimeters) were −45.1, −8.9, and 1.9 in the left STG, 43.1, −2.6, and 2 in the right STG, −40.8, 18.5, and 15.6 in the left IFG, and 37.6, 21.2, and 15.1 in the right IFG, respectively (Fig. S1A). The signal for ECDs was extracted in 400 ms epochs after the onset of the ending chord using 1–20 Hz band-pass filter for each participant. The 400 ms was the time window involving the peak latencies of P2m and ERANm in our previous studies (Kim et al., 2011; Kim et al., 2014) (Fig. S1B).

Using the ECDs signals of the time window of 400 ms in the bilateral IFGs and STGs for each condition, we estimated the information flows in 12 directional connections between the bilateral IFGs and STGs for the three conditions. Effective connectivity for 12 connections was calculated by LTDMI (Jin et al., 2010). The LTDMI is an information theoretic measure of functional coupling based on mutual information (MI) (Jin et al., 2011; Jin et al., 2012) which predicts information transmission between two time series.

MI is defined as the quantity of information shared in two time series of *X*(*n*) and *Y*(*n*) (*n* = 1, 2,…, N), at N discrete points. The probability density function (PDF) of *X*(*n*) and*Y*(*n*) are *p*(*X*(*n*),*κ*) ≡ *p*(*X*(*n*)) and *p*(*Y*(*n*),*κ*) ≡ *p*(*Y*(*n*)) with *n* = 1, 2,…, bin, respectively. The MI is computed by *p*(*X*(*n*), *Y*(*n*)), the joint PDF between *X*(*n*) and *Y*(*n*), as follows:

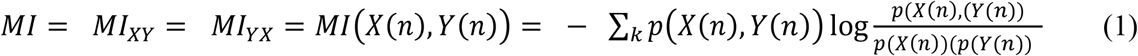

If *X*(*n*) and *Y*(*n*) are completely identical, the MI is maximum. However, if two time series are independent of each other, the MI is zero. The directional information transmission between the two time series can be calculated by time delayed mutual information (TDMI) (Jin et al., 2010):

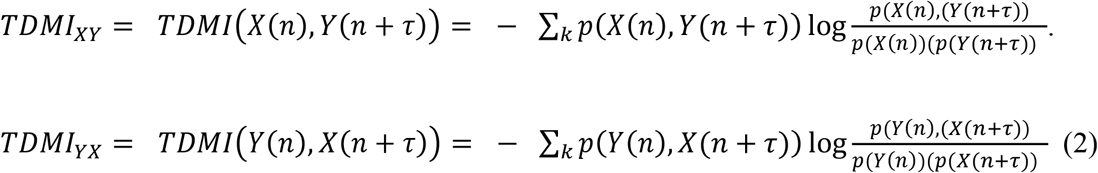

TDMI can detect linear and non-linear correlation between two time series. Since the data length used in the present study (400 ms epoch) was insufficient to reconstruct a reliable PDF for general TDMI presented in equation (2), we used LTDMI as an effective connectivity measure in this study. LTDMI is adopted as follows:

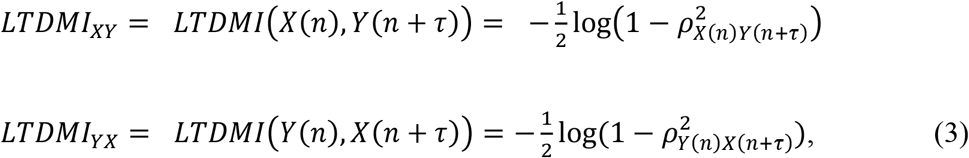

where *ρ*_*X*(*n*)*Y*(*n*+*τ*)_ and *ρ*_*Y*(*n*)*X*(*n*+*τ*)_ are a cross-correlation coefficient, and τ is delay time, which was 120 ms in our present study. To estimate the linearized information flow between the time series, each time series is assumed with the Gaussian distributed function with zero-mean, and variance 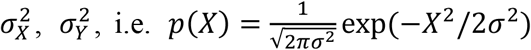. The LTDMI values were averaged over delay time.

Differences in the LTDMI values among the three conditions of T, SM, and ST in 12 connections were tested by the two-way repeated measures ANOVA. In all *post hoc* analysis steps, the alpha levels for multiple comparisons were adjusted by the FDR correction (*P* < 0.05). Additionally, the group difference for the LTDMI values was tested by the independent *t* test (*P* < 0.05).

In the MEG experiment, the mean CR for staccato chord detection was calculated for each participant. In the behavioral experiment, the difference between the three conditions for the CR was determined by the one-way repeated measure for all conditions.

Correlation analysis was performed to test the relationships between the LTDMI value and CR for all conditions of all participants (i.e., for the merged data set of 3 conditions of 18 participants). In the correlation analysis for the LTDMI value and the CR, the correlation was tested using the Spearman rank correlation because the data was not normally distributed. The correlation was calculated using the one-tailed test, because the ambiguous stimuli lead to slower and less accurate responses than the easy stimuli (Sabri et al., 2006; Jastorff et al., 2009; Fleming et al., 2010). The alpha level was adjusted by the Benjamin-Hochberg false discovery rate (FDR) correction for the multiple comparisons testing of the three conditions (*P* < 0.05). The Greenhouse-Geisser’s correction was applied because the sphericity of the data was violated via the Mauchly sphericity test. All statistical analyses were performed using SPSS 21.0 software (IBM, Armonk, NY, USA).

## Results

### LTDMI values for three conditions

Using the LTDMI (Jin et al., 2010), we calculated effective connectivity for 12 connections among 4 regions of interest (ROIs) of the bilateral IFGs and STGs for three conditions of the T, SM, and ST in 19 participants. For the LTDMI values, we performed a two-way repeated measures analysis of variance (ANOVA) with two factors of Condition and Connection. The ANOVA (n = 19) showed a significant main effect of Condition [*F*(1.872, 404.247) = 3.108, *P* = 0.049] and a significant interaction of Condition × Connection [*F*(20.587, 404.247) = 2.555, *P* = 0.0002], and a significant effect of Connection [*F*(1, 216) = 1.920, *P* = 0.038]. *Post hoc* one-way repeated measures ANOVAs with the Condition factor in 12 connections confirmed a connection reflecting the difference among the three conditions. The difference between the three conditions was revealed only in two connections from the right to the left IFG [*F*(2, 36) = 6.526, *P* = 0.024, false discovery rate (FDR) corrected] and from the right to the left STG [*F*(2, 36) = 12.373, *P* < 0.001, FDR corrected] among 12 connections (Fig. 2B,C; see also Table S1).

**Figure 2.**
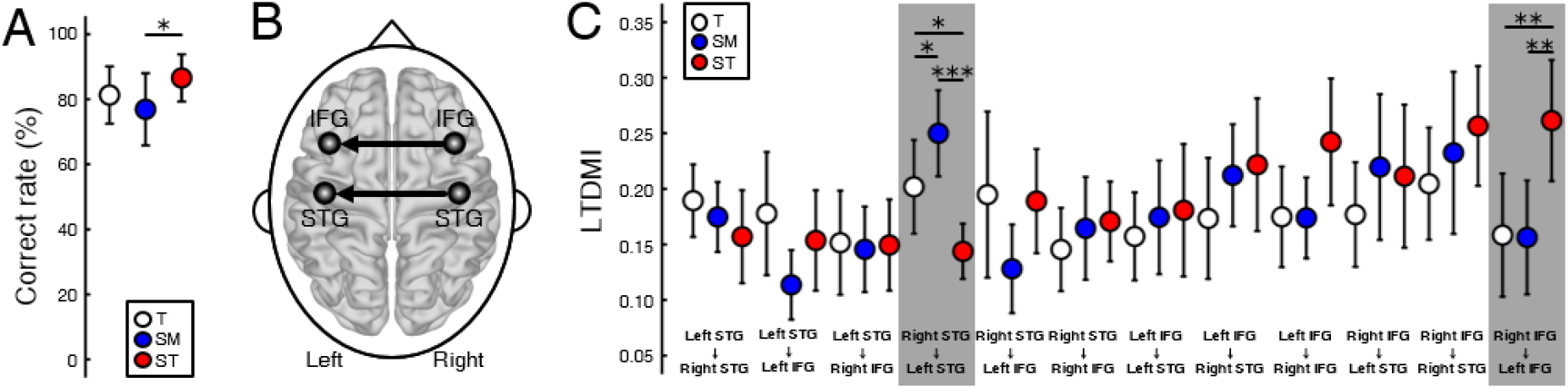
Difference in the LTDMI values for the three conditions. (A) CF of SM is significantly lower than the CR of ST. The other pairs are not statistically significant. * *P* < 0.05. (B) Difference between the conditions for LTDMI was revealed in only two interhemispheric connections, which were termed “IFG-LTDMI” and “STG-LTDMI”. (C) The STG-LTDMIs were different between all pairs. In the IFG-LTDMI, the SM was higher in the other conditions. * *P* < 0.05, ** *P* < 0.01, and *** *P* < 0.001 (FDR corrected). Error bars denote 95% confidence intervals. See also Table S1.

Hereafter, we use the term “IFG-LTDMI” to refer to the LTDMI values from the right to the left IFG, and the term “STG-LTDMI” to refer to those from the right to the left STG. In the two interhemispheric connections, the SM and ST of the most ambiguous and irregular conditions showed the highest STG-LTDMI and IFG-LTDMI, respectively. In a *post hoc* paired *t* test, the IFG-LTDMI was higher for the ST than for the T [*t*(17) = 3.289, *P* = 0.009, FDR corrected] and for the SM [*t*(17) = 2.711, *P* = 0.009, FDR corrected], while the IFG-LTDMI for the T and SM was not significantly different [*t*(17) = −0.061, *P* = 0.952, FDR corrected]. The STG-LTDMI was higher for the SM than for the T [*t*(17) = −2.691, *P* = 0.023, FDR corrected], and the ST [*t*(17) = 5.357, *P* = 0.0001, FDR corrected], while was significantly higher for the T than for the ST [*t*(17) = 2.259, *P* = 0.037, FDR corrected].

### Behavioral response

During magnetoencephalography (MEG) experiment, participants were asked to listen to each condition carefully and to detect the sequences including a staccato chord in order to check the level of attending to the condition (Fig. 1B). All participants detected the staccato chord with more than 95% including the number of missed buttons. This indicates that the participants paid attention to musical stimuli. After the MEG experiment, participants performed a behavioral test discriminating among the three conditions (Fig. 1B). The mean CR (n = 18) was lower in the SM (77.0%) than in the T (82.4%) and the ST (88.7%). The one-way repeated measure ANOVA (n = 18, excluded 1 outlier) showed a significant main effect of Condition [*F*(2,34) = 4.799, *P* = 0.015]. In a *post hoc* analysis, the difference between the CRs was significant only between the SM and the ST (Fig. 2A). The SM was significantly lower than the ST [*t*(17) = −2.574, *P* = 0.020]. There were no significant differences in the pairs of T vs. SM [*t*(17) = 1.772, *P* = 0.094] and T vs. ST [*t*(17) = −1.753, *P* = 0.098].

### Correlation between the LTDMI values and correct rate

To confirm whether the STG-LTDMI reflected perceptual ambiguity, and, if it was so, whether it was specific for the STG-LTDMI among the IFG-LTDMI and the STG-LTDMI, we tested the correlation between the LTDMI values of STG-LTDMI/IFG-LTDMI and the behavioral response of CR. The correlation was tested using the values for all conditions and participants (*n* = 57, 3 conditions × 19 participants). A significant correlation with the CR was observed not in the IFG-LTDMI but in the STG-LTDMI (one-tailed Spearman’s rank correlation; STG-LTDMI, Spearman’s *ρ* = −0.260, *P* = 0.026; IFG-LTDMI, Spearman’s *ρ* = 0.064, *P* = 0.319) (Fig. 3).

**Figure 3.**
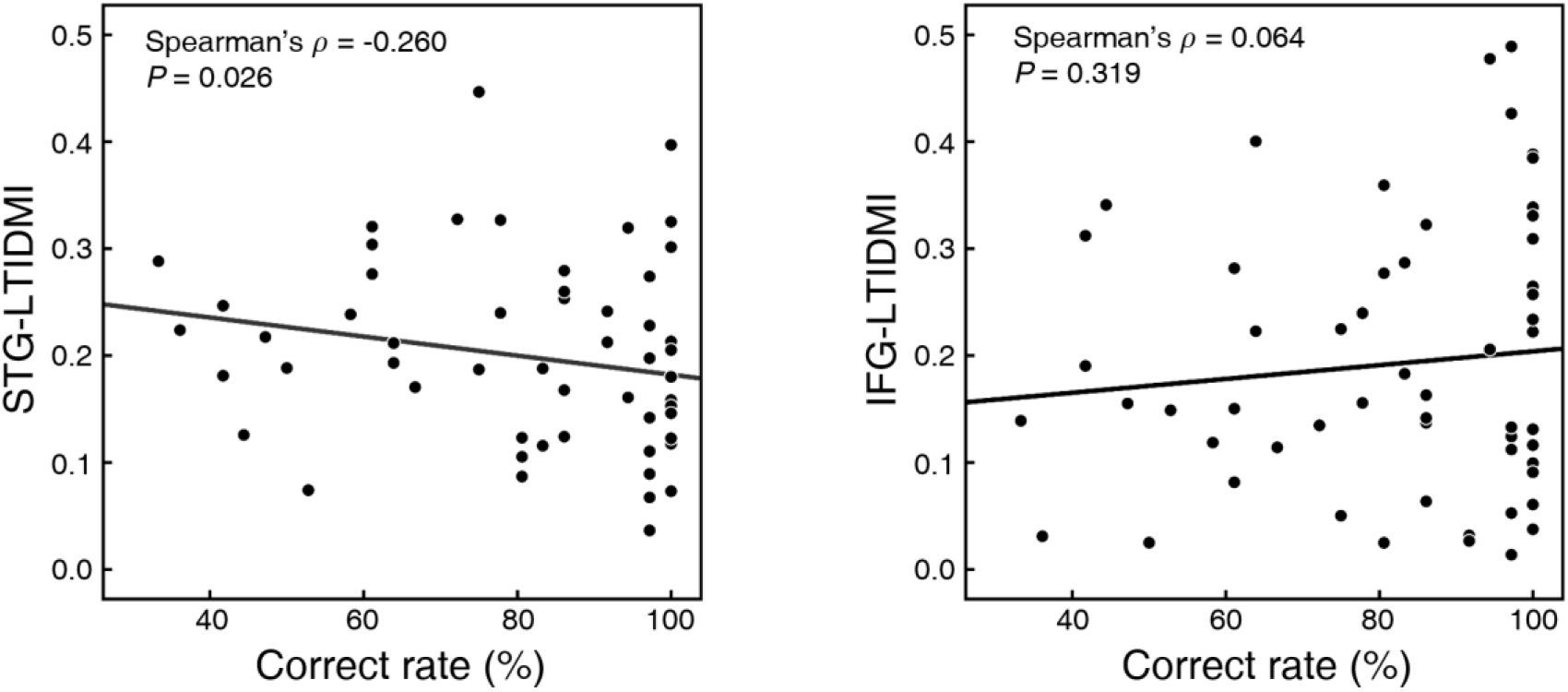
Correlation between CR and STG-LTDMI/IFG-LTDMI. The CR was only negatively correlated with the STG-LTDMI (*n* = 57, one-tailed Spearman’s rank correlation). However, the CR did not show a correlation with the IFG-LTDMI (*n* = 57, one-tailed Spearman’s rank correlation).

## Discussion

The IFG-LTDMI was enhanced for the ST of the most irregular condition. The STG-LTDMI was enhanced for the SM of the most ambiguous condition. The processing of syntactic irregularity and perceptual ambiguity in the three conditions was dissociated in the IFG-LTDMI and the STG-LTDMI, respectively. This implies that the brain interprets the three conditions as both “from regular to irregular” and “from ambiguous to unambiguous” conditions simultaneously.

The highest IFG-LTDMI for the ST is a further extension of the highest ERAN response elicited only for the irregular condition (ST) in our previous study, which found that a prominent early right anterior negativity’s magnetic counterpart (ERANm) was not observed in the SM of the less irregular condition in terms of syntactic irregularity (Kim et al., 2014). The IFG-LTDMI in terms of effective connectivity may underlie the ERAN. Moreover, the IFG-LTDMI from the right to the left IFGs is one step forward from the previous reports that the bilateral IFGs are the neural generator of ERAN (Maess et al., 2001; Sammler et al., 2009; Kim et al., 2011; Kim et al., 2014). Our data suggests that the left IFG and the right IFG interrelate in the processing of musical syntax, in terms of effective connectivity.

The STG-LTDMI was highest for the most ambiguous SM. The ST was lowest. Unambiguous stimuli of speech and ambiguous stimuli of speech like song differently activate the STG (Tierney et al., 2013). The patterns of STG-LTDMI between all conditions were consistent with our hypotheses that the SM would be most ambiguous. Moreover, these results show that the T was more ambiguous than the ST with the salient highest voice. The smallest unit comprising of harmony and melodic line in chord sequence is a single tone. In language, a phoneme is the smallest unit. Acoustic–phonetic processing is related with the STG (Callan et al., 2004). Auditory areas involving Heschl’s gyrus are activated by ambiguous phonems (Kilian-Hutten et al., 2011). Based on the aforementioned studies, we interpret our findings in the bilateral STGs as indicating neural substrates for the perception of ambiguity implied in harmony and melodic line in chord sequence.

In previous studies, the connection between the IFG and STG is involved in the processing of syntax in music and language (Sakai et al., 2002; Sammler et al., 2009; Friederici, 2011; Musso et al., 2015). Moreover, a fMRI study using a real musical piece reported that the different level of syntactic irregularity was reflected in functional connectivity between the IFG and STG (Seger et al., 2013). In another fMRI study on acoustic–phonetic processing, IFG-STG coupling is increased by ambiguous acoustic signal (Leitman et al., 2010). However, our data did not show the connection between the IFG and the STG in either syntactic irregularity or perceptual ambiguity. Instead, the connectivity was dissociated in each of IFG and STG. Furthermore, the direction of effective connectivity was from the right to the left hemisphere in both the IFG-LTDMI and STG-LTDMI. We interpret dissociation of the IFG and STG in connectivity as indicating the functional segregation related with syntax and ambiguity processing. Also, we interpret that the same direction of information transmission in the bilateral IFGs and STGs as indicating the different roles of bilateral hemispheres in music processing and indicating the IFG-LTDMI and STG-LTDMI commonly based on the rightward asymmetry.

Considering the STG-LTDMI, the SM is most ambiguous among the three conditions, and the ST is less ambiguous than the T. The more ambiguous condition may be more difficult to response than the less ambiguous condition. However, the CR was only significantly different between the SM and the ST. Our results demonstrate that the ST with a note of “D” in the highest voice, not included in the other conditions, was most unambiguous. These are consistent with previous studies addressing the perceptual prominence of melodic line (Marie and Trainor, 2013). Music-majors were more sensitive to harmony, while non-music-majors were more sensitive to melodic line (Fujioka et al., 2005). Our results show that the relationship between harmony and voice leading importantly affects chord perception.

Furthermore, for all participants, a significant correlation was only observed in the STG-LTDMI, not in the IFG-LTDMI. Also the following point should be taken into account. In the MEG experiment, the participants were asked to detect the “staccato” chords. Thus, both IFG-LTDMI and STG-LTDMI, based on the MEG data, would not reflect the attentive processing of the three conditions. Syntactic irregularity for harmony and melody of the highest voice can be perceived pre-attentively, which elicits an ERAN response (Koelsch et al., 2002b). Moreover, a deviant tone in tone sequence, eliciting mismatch negativity (MMN), can also be processed unconsciously (Brattico et al., 2006; Durschmid et al., 2016). This indicates that both syntactic irregularity and perceptual ambiguity based on harmony and melodic line can be perceived pre-attentively. Nevertheless, there was a positive correlation between STG-LTDMI and CR, although the STG-LTDMI was pre-attentive and the behavioral CR was attentive.

Though our data suggest perceptual segregation of syntactic irregularity and perceptual ambiguity in chord sequences, we could not conclude that the present findings are relevant to whole chords in Western music theory. This would be completed by using musical stimuli specifying the level of syntactic irregularity and perceptual ambiguity. Moreover, our data only focused on the 4 ROIs of the bilateral IFGs and STGs based on our hypothesis. The syntactic irregularity and perceptual ambiguity processes responding to the whole chord of Western tonal music would be also examined in terms of the relevant brain regions including the 4 ROIs of the bilateral IFGs and STGs on our hypothesis.

Nevertheless, we demonstrate for the first time that interhemispheric connectivity in the bilateral IFGs and STGs respectively dissociates syntactic irregularity and perceptual ambiguity in chord sequences. Our results suggest that syntactic irregularity and perceptual ambiguity in music, are processed simultaneously and separately.

## Author Contribution

C.H.K and C.K.C. conceived the study. S.H.J. and J.S.K. contributed analytic tools. Y.K. and S.W.Y. contributed music-theoretical background. C.H.K. analyzed the data. C.H.K. and C.K.C. wrote the paper. All authors discussed the results and reviewed the paper.

## Acknowledgements

We would like to thank Ji Hyang Nam for the technical support in MEG acquisition.

