## Supplementary Figure 1 and Supplementary Table 1 for "Dissociation of connectivity for syntactic irregularity and perceptual ambiguity in musical chord stimuli"

### Supplementary information

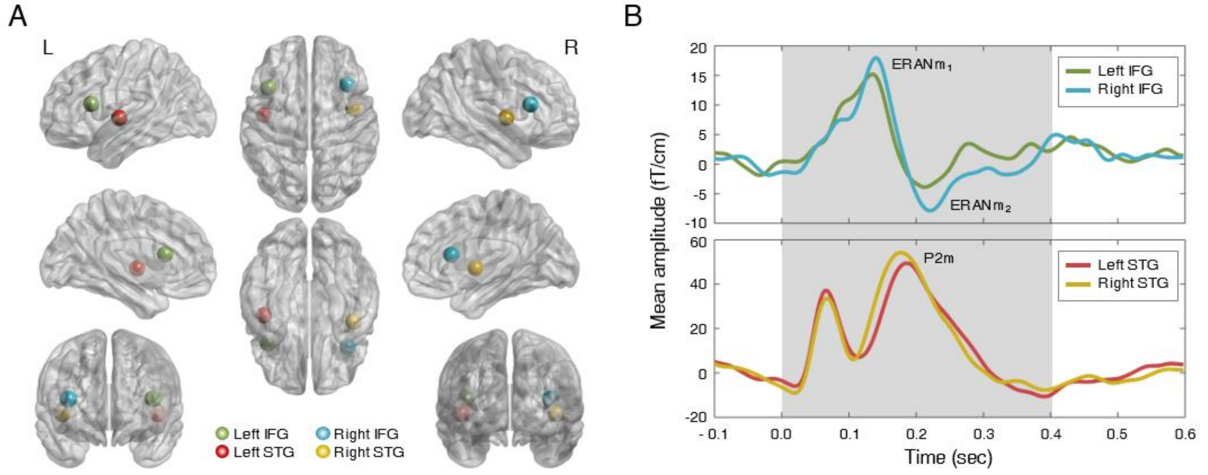


**Figure S1.** Mean dipole source locations of all participants in the bilateral IFGs and STGs, and the time window. (A) The ECDs of P2m (magnetic counterpart of P2) and ERANm (magnetic counterpart of ERAN) in the ending chord in each condition were localized in the bilateral IFGs and STGs. The green dot (left IFG), sky blue dot (right IFG), red dot (left STG), and yellow dot (right STG) indicate the grand average (*n* = 19) of individual ECDs estimated by BESA 5.1.8.10. (x, y, and *z in* Talairach coordinates, millimeters, left IFG, -40.8, 18.5, and 15.6; right IFG, 37.6, 21.2, and 15.1; left STG, -45.1, -8.9, and 1.9; right STG, 43.1, -2.6, and 2). The brain map and dipole source location were generated by BrainNet Viewer (http://nitrc.org/projects/bnv/). (B) The LTDMI values were calculated in the time window of 0 ms to 400 ms after the ending chord onset in the MEG signal of ECDs for four ROIs. The green, sky blue, red, and yellow lines denotes the mean amplitudes of three conditions (*n* = 19) in each ROI across all participants. The gray shadow indicates the time window of 400 ms encompassing the peak latencies of $ERANm_1$ and $ERANm_2$ in the bilateral IFGs and P2m in the bilateral STGs, respectively.

**Table S1.** *Post hoc* one-way repeated measures ANOVAs results of the LTDMI for the Condition factor in 12 connections. The *P*-values were corrected by FDR for multiple comparisons of 12 connections. *$P < 0.05$, and ***$P < 0.01$.

| | *df* | *F* | *P* |
|---|---|---|---|
| ***Left STG → Right STG*** | 2/36 | 1.275 | 0.501 |
| ***Left STG → Left IFG*** | 2/36 | 3.235 | 0.153 |
| ***Left STG → Right IFG*** | 2/36 | 0.023 | 0.977 |
| ***Right STG → Left STG*** | 2/36 | 12.373 | ***** < 0.001** |
| ***Right STG → Left IFG*** | 2/36 | 4.828 | 0.056 |
| ***Right STG → Right IFG*** | 2/36 | 0.478 | 0.749 |
| ***Left IFG → Left STG*** | 2/36 | 0.234 | 0.864 |
| ***Left IFG → Right STG*** | 1.399/25.187 | 1.024 | 0.522 |
| ***Left IFG → Right IFG*** | 1.220/21.951 | 3.030 | 0.178 |
| ***Right IFG → Left STG*** | 2/36 | 0.905 | 0.552 |
| ***Right IFG → Right STG*** | 2/36 | 2.855 | 0.170 |
| ***Right IFG → Left IFG*** | 2/36 | 6.526 | *** 0.024** |
